# Color polymorphism in the saddleback clownfish, *Amphiprion polymnus*: species or complex?

**DOI:** 10.64898/2026.06.28.735040

**Authors:** Lucy M. Fitzgerald, Floriane Coulmance, Anna Marcionetti, Théo Gaboriau, Alberto García Jiménez, Prince T. Apag, Melissa Versteeg, Fletcher J. Noble, Kirstin Gaffney, Manon Mercader, Annie G. Diola, Paul John Geraldino, Theresa Rueger, Vincent Laudet, Nicolas Salamin

## Abstract

Color polymorphism can facilitate local adaptation, maintain intraspecific diversity, or reflect early stages of speciation. Clownfishes (*Amphiprion spp*.) typically display a simple black, orange, and white pattern, but the saddleback clownfish (*Amphiprion polymnus*) shows striking variation in melanism and the number of vertical bars, which are thought to play a role in species recognition. In 2024, a revision on iNaturalist split *A. polymnus* into multiple species based solely on color pattern and geographic range. This raises the question of whether these morphs represent true species or intraspecific polymorphism, which we tested using genomic and image-based data. We sampled 97 individuals from seven populations across the species’ range and quantified color patterns from standardized photographs. Phenotypic and genomic analyses reveal a complex pattern of divergence. Image analysis identified three distinct phenotypic clusters, with *A. polymnus, A. annamensis*, and *A. laticlavius* each showing consistent differences in saddle shape and vertical bar extent. ADMIXTURE resolved three distinct genetic groups corresponding to the morphs. Pairwise *F*_*ST*_ (0.54–0.71) and *d*_*xy*_ indicate extremely high differentiation between *A. polymnus* and *A. annamensis*, consistent with species-level divergence, whereas *A. laticlavius* shows much lower differentiation from *A. polymnus* (*F*_*ST*_ 0.09–0.18) and higher differentiation from *A. annamensis* (*F*_*ST*_ 0.64–0.66). Overall, phenotypic and genomic data show structured variation, but the status of *A. laticlavius* remains ambiguous. Our study reveals clear and structured divergence across the full range, yet the taxonomic interpretation of this variation remains inherently challenging. The key question remains: do these patterns reflect a single polymorphic species or a complex of closely related species?

## Introduction

Color polymorphism plays a central role in evolutionary processes, generating phenotypic diversity both within and between species. Over time, this variation can either be maintained, lost, or give rise to new speciation (Kusche et al., 2015). One of the most well-known study systems for color polymorphism is the adaptive radiation of African cichlid fish (Seehausen, 2006; Seehausen et al., 1999). In this group, variation in coloration is closely associated with ecological specialization, such as habitat use (e.g. rocky substrate vs. vegetative environments). This has played a key role in both niche partitioning and reproductive isolation. These examples highlight how polymorphic traits can structure populations and promote diversification.

In some cases, persistent polymorphism may represent an early stage of divergence, blurring the boundary between within-species variation and the formation of species complexes. Species complexes consist of closely related lineages with different ecological niches and genetic signatures of adaption based on their niches. In the marine environment, a well-studied example is the *Dascyllus trimaculatus* complex, in which multiple lineages display distinct phenotypes despite limited morphological divergence (Roberts, 2022). Interestingly, this group represents one of the few Pomacentridae alongside the clownfish (*Amphiprion*) group that associate with sea anemones during their juvenile stage.

Clownfishes *Amphiprioninae* are a monophyletic group of 28 species that live in an mutualistic association with sea anemones that likely triggered their adaptive radiation (Litsios et al., 2012). Coloration in clownfish has been linked to their ecological specialization, such as their anemone host. For example, species specializing on *Radianthus* sea anemones are more orange/yellow, whereas *Entacmaea* specialists are dark orange/red. Generalists, which can inhabit more than three species of sea anemones are darker with contrasting vertical bars (Gaboriau et al., 2025). This suggests that coloration plays a role in ecological adaptation within the group.

Within clownfishes, there are two species *Amphiprion clarkii* and *A. polymnus*, that are considered to be polymorphic in the number of white vertical bars Salis et al. (2018). This has been hypothesized to function as species recognition, as cohabitation between species is rare, but when it occurs, they never have the same number of vertical bars (Camp et al., 2016). Both species also display behavior optimized for swimming and spends most of its time outside the anemone (Mercader et al., 2025), which may influence the evolution of coloration. Despite this shared polymorphism, their relationship between coloration and population structure differs between the species. In *A. clarkii*, which has the widest geographic range and is considered the most generalist clownfish inhabiting all 10 sea anemone hosts, coloration was not linked to the population structure. Instead, the highest variability in color variation was at the center of *A. clarkii*’s distribution and the edge of the species range did not overlap in color pattern (Schmid et al., 2024). However, recent work (Zwahlen et al., in prep) suggests that *A. clarkii* may be a species complex with *A. tricinctus*, its sister species.

In contrast, the saddle-back clownfish, *A. polymnus* is a specialist found primarily in *Stichodactyla* anemone species although recently has been found to be living in *Macro doorensis* (Astakhov et al., 2025). *A. polymnus* lives in sandy or mucky areas that are lesser known diving spots and therefore this species remains relatively understudied. *A. polymnus* is part of the Polymnus clade along with its sister species, *A. sebae* which can also live in sandy/mucky areas. Body-wise, it differs from other clownfish clades with an oval shape (Allen, 1980). Color-wise, the body is primarily black with a single white vertical bar on the head and a white saddle on the upper dorsal fin. The width of the saddle as well as a third vertical bar on the peduncle can vary greatly in shape and color (Fautin & Allen, 1992). In Fautin, D. G. (1991) described that orange portions of clownfish species such as *A. polymnus* can darken when they occupy anemone host *Radianthus* (formerly *Heteractis crispa*). Yet, no further description of this color change has been mentioned since.

Recent taxonomic uncertainty further complicates the interpretation of this color variation. In October 2024 on iNaturalist, the species distribution changed and resurrected a previously synonymous name of *A. polymnus*, called *A. annamensis* which spans the distribution of the South China Sea from Thailand to the Philippines, North to Japan and Taiwan citing CoF and amphiprionology.wordpress.com. The blog describes *A. annamensis* as having a broad middle stripe that does not reach the belly and no posterior stripe. Additionally, the body and fins are highly variable from solidly orange, to solid black. The type locality is from the south coast of Vietnam. The “true” *Amphiprion polymnus* is found in Indonesia, from Java Sea and Bali east to Raja Ampat and north to Southern Mindanao and rarely reported from Cebu and Samal in the Philippines. The caudal fin is black with white margins and the middle vertical bar reaches the belly a posterior vertical bar is present. The type locality is “Indiis”, which is presumed to be Indonesia. The name *Amphiprion bifasciatus annamensis* was first described by Chevey in 1932 but became synonymous for *Amphiprion polymnus* (Linnaeus, 1758) by G.R. Allen in 1991. The blog also describes the Polymnus complex which includes *A. polymnus, A. laticlavius, A. annamensis* and *A. sebae* of which only two of the species are recognized (*A. polymnus* and *A. sebae* as sister species). *A. laticlavius* (Cuvier 1930) is similar to the *A. annamensis* which has a similar middle stripe however in this species, the middle stripe is typically more thickened and the base color is more orange-brown than yellow orange and the melanism is more variable in extent and tends to form first medially rather than dorsally. The type locality is New Guinea with a range from Melanesia, from Raja Ampat to the Solomon Islands. The blog writes that for *A. annamensis* and *A. laticlavius*, “This species was treated as a synonym of *Amphiprion polymnus* in Allen 1991 and subsequent references. The elevation to a full species status used in this classification should be considered provisional until a full taxonomic revision is published.”

Currently, there are 10 synonyms of *A. polymnus* on WoRMs and CoF, and on both sites *A. annamensis* has been validated as its own species. However, the reference for elevating *A. annamensis* has been recent (2024) with no scientific publication or additional evidence presented CoF. What is currently recognized as *A. polymnus* may represent a species complex of multiple distinct lineages.

The question remains, does the observed variation in coloration within *A. polymnus* reflect true polymorphism within a single species or does it represent a species complex that were previously described species and then combined into a single name by a single author? Resolving this distinction is not only important for understanding the evolutionary processes shaping color variation, but also for conservation, as species boundaries influence management strategies. This paper aims to disentangle the genomic basis of color variation in *A. polymnus* to test whether it represents a true color polymorphism or evidence of ongoing divergence within a species complex. We hypothesize that genomic population structure is associated with coloration patterns, and we use whole-genome sequencing and quantitative color analysis to evaluate whether distinct morphs correspond to genetically differentiated lineages.

## Methods

### Data Collection

We analyzed *Amphiprion polymnus* samples from five countries: Japan, Thailand, Philippines, Indonesia, and Papua New Guinea across seven geographical regions: Seragaki Bay (JPN), Koh Tao/Koh Phangan (THA), Dauin/Panglao as Negros/Bohol (NEG/BOH), Mindanao (MIN), Sulawesi (SUL), Bali (BAL), and Kimbe Bay (PNG). The geographic regions were determined first by country, and then by island if there was more than one location sampled within the countries. For Thailand Koh Tao and Koh Phangan were morphed into one location as well as the two islands in the central Philippines (Bohol and Negros Oriental) (Figure 1A). The samples were collected during SCUBA diving expeditions. We gathered genomic (fin clip), ecological, and phenotypic (JPEG photographs) data for the entire dataset (n = 97). For 51 individuals, (Philippines and a subset of the Papua New Guinea dataset) we obtained standardized photos (RAW photographs) using a underwater frame that places the camera at a fix distance and angle from a pvc background with a neutral color and XRite color checker. Previously published genomic data from García-Jiménez et al., 2025 was added for Thailand (n = 10) and Indonesian samples (n = 8). New samples (n = 79) were added from Japan (n = 14), Philippines (n = 41), and Papua New Guinea (n = 24), for a total of 97 genomic samples. Samples were preserved either in 70% ethanol (EtOH) or DESS buffer (Oosting et al., 2020).

**Figure 1:**
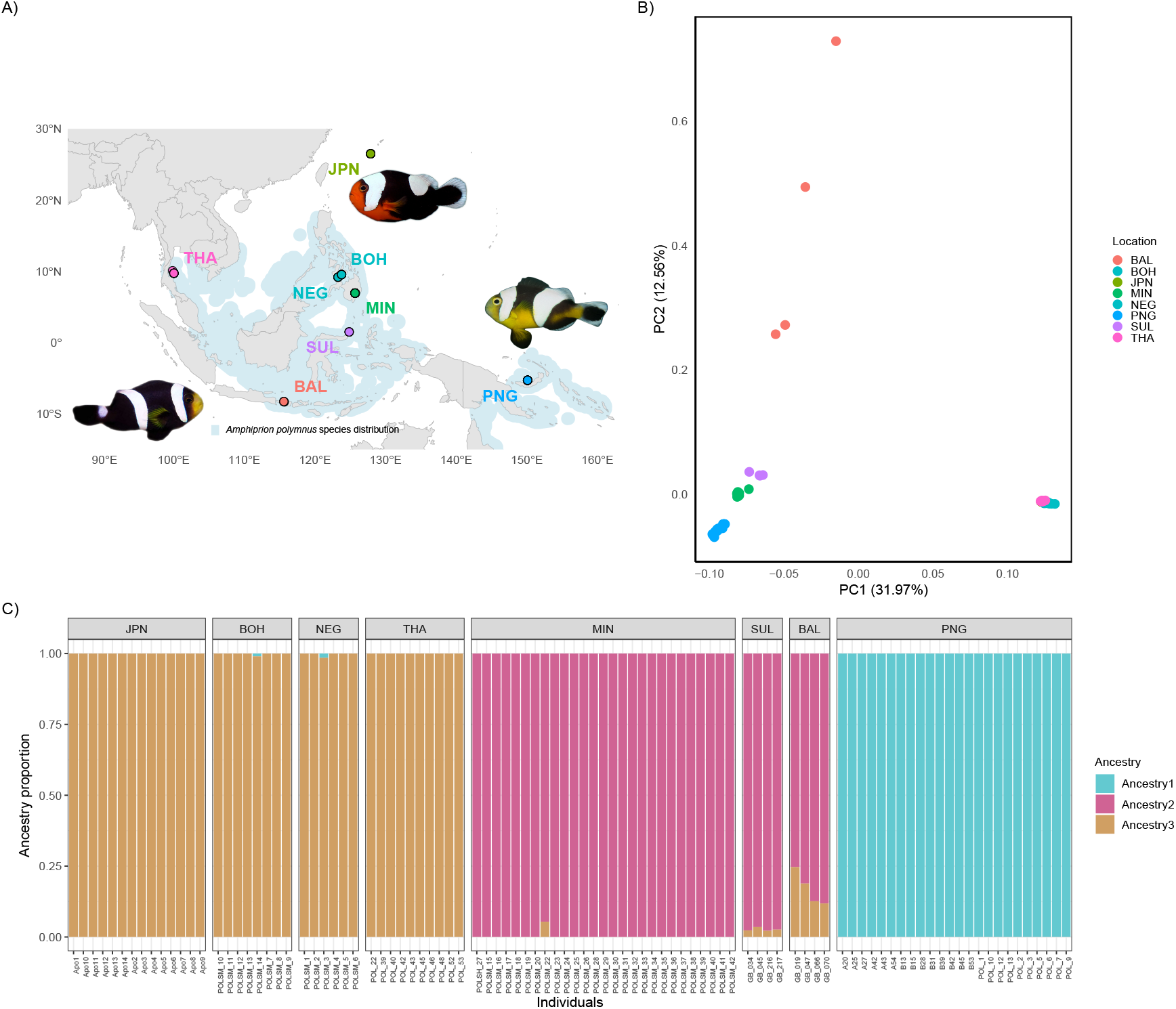
A) The map of sampling locations, each dot represents a location colored by region: Japan (JPN) in light green, Bohol (BOH) in cyan, Negros Oriental (NEG) in cyan, Thailand (THA) in pink, Mindanao (MIN) in dark green, Sulawesi (SUL) in purple, Bali (BAL) in salmon, and Papua New Guinea (PNG) in blue with the three morphs, *A. annamensis* in the north (JPN, BOH, NEG, and THA), *A. laticlavius* to the east (PNG), and the true *A. polymnus* to the south (BAL, SUL, and MIN) (relative to the center of the distribution) B) Genomic PCA of SNPs (1,058,491 SNPs) showing 2 clusters, BAL, SUL, MIN, and PNG vs. BOH, JPN, NEG, and THA C) ADMIXTURE analysis found the best *K* = 3 with three ancestries: 1) JPN, BOH, NEG, and THA 2) MIN, SUL, and BAL and 3) PNG.

### DNA extraction, library preparation and sequencing

DNA was extracted using DNeasy blood and Tissue Kit (Qiagen GmbH) and quantified using Qubit^®^ 2.0 Fluorometer (Thermo Fisher Scientific). DNA libraries were prepared using the Nextera DNA Flex Library Preparation Kit following the manufacturer’s instructions. The fragment length distribution of the libraries was validated using a Bioanalyzer (Agilent Technologies). Libraries were sequenced in two runs: either on the NovaSeq 6000 Sequencing System (Illumina) or Aviti (Element Biosciences) with 150 paired-end lanes. Sequencing was performed at the Genomic Technologies Facility (GTF) of the University of Lausanne, Switzerland.

### Reads processing, mapping and SNP calling

Raw reads were trimmed and adapters were removed using *Trimmomatic* v.0.39 (Bolger et al., 2014). The quality was checked before and after trimming using (*FastQC* v.0.11.9) and *MultiQC* (Ewels et al., 2016). Reads were mapped against a high quality PacBio genome, *A. polymnus* (Marcionetti et al., in prep) using *BWA* v0.7.17 (Li & Durbin, 2009); and *Samtools* v1.19.2 (Danecek et al., 2021) was used to process and sort the mapped reads, generating mapping statistics with *Bamtools* v2.5.2 (Barnett et al., 2011).

Haplotypes were called with *GATK* v4.5.0.0 (Van der Auwera & O’Connor, 2020), then the gVCF files were merged with (*Picard Tools* v3.1.1) and proceeded to joint-genotyping with *GATK*. We used *VCFtools* v0.1.16 and kept only biallelic SNPs with quality above 30 (`-minQ 30`), minimum depth 5 (`-minDP 5`), maximum depth 40 (`-maxDP 40`), max missingness 0.95 (`-max-missing 0.95`) and a minor allele frequency (`-MAF 0.01`).

### Population Genomics Analyses

PCA & ADMIXTURE: We selected independent SNPs by performing linkage disequilibrium (LD) pruning using *PLINK* v1.9 (Purcell et al., 2007) (`--indep-pairwise 50 10 0.1`). Then performed a PCA and estimated the individual admixture proportions using *ADMIXTURE* v1.3.0 (Alexander et al., 2009) with *K* -values ranging from 2-5. The *K* -value with the lowest cross-validation error was chosen (K = 3).

*F*_*ST*_ : We also investigated divergence and diversity among the different populations (geographic regions defined in Figure 1) by estimating between-population differentiation (*F*_*ST*_), between-population absolute divergence (*d*_*xy*_) as well as population nucleotide diversity (*π*) in non-overlapping 5000 SNPs windows using a custom script popgenWindows.py.

### Image Analysis

The standardized image analysis dataset consisted of 51 standardized photographs.Photographs of individual clownfish were taken in situ, using a underwater frame that places the camera at a fix distance and angle from a PVC background with a neutral color and XRite color checker. Fish is captured in its anemone, transferred to the PVC frame for photographing, a fin clip is taken and then the fish is released on the same anemone as it was taken from. We processed raw photographs following the pipeline described in Coulmance et al., 2024. We performed a PCA analysis to capture the color pattern variation. Significance was tested with a PERMANOVA.

## Results

### Population Genomics Analyses

We used a PCA on the covariance matrix to examine the population structure of *A. polymnus*. On the first PC axis (explaining 31.97% of the variation) we found a clear separation between BOH, NEG, JPN, and THA vs. the rest of the populations corresponding to the separation of the morphs *A. annamensis* from *A. polymnus* and *A. laticlavius*. On the second PC axis (explaining 12.56% of the variance), we see a split between PNG, MIN, SUL vs. BAL although this could be driven by the low sample size (n=4) (Figure 1B).

To examine ancestral introgression among populations, we used ADMIXTURE to calculate the admixed proportion for each individual. Using the cross validation, the best K = 3 splitting the populations into 3 groups corresponding to the three morphs: 1) JPN, BOH, NEG, and THA (*A. annamensis*) 2) MIN, SUL, and BAL(*A. polymnus*) and 3) PNG (*A. laticlavius*). The first and third cluster had almost pure ancestry whereas the second cluster had some admixed individuals particularly around BAL suggesting that currents play a role in the gene flow between the populations (Figure 1C).

We then examined patterns of genomic difference, divergence and diversity by calculating *F*_*ST*_ and *d*_*xy*_ between each pair of populations and *π* within populations in 5000 SNPs windows. We observed high *F*_*ST*_ values ranging from 0.014 (SUL vs. MIN) to 0.713 (SUL vs. THA). We found a similar pattern in *d*_*xy*_ with means ranging from 0.019 (JPN vs. THA) to 0.127 (JPN vs. PNG). Both the *F*_*ST*_ and *d*_*xy*_ patterns show low gene flow between *A. annamensis* from *A. polymnus* and *A. laticlavius* but the intermediate gene flow between *A. polymnus* and *A. laticlavius* is less clear, similar to the genomic PCA. *π* values ranged from 0.010 (THA) to 0.045 (BAL) (Figure 2).

**Figure 2:**
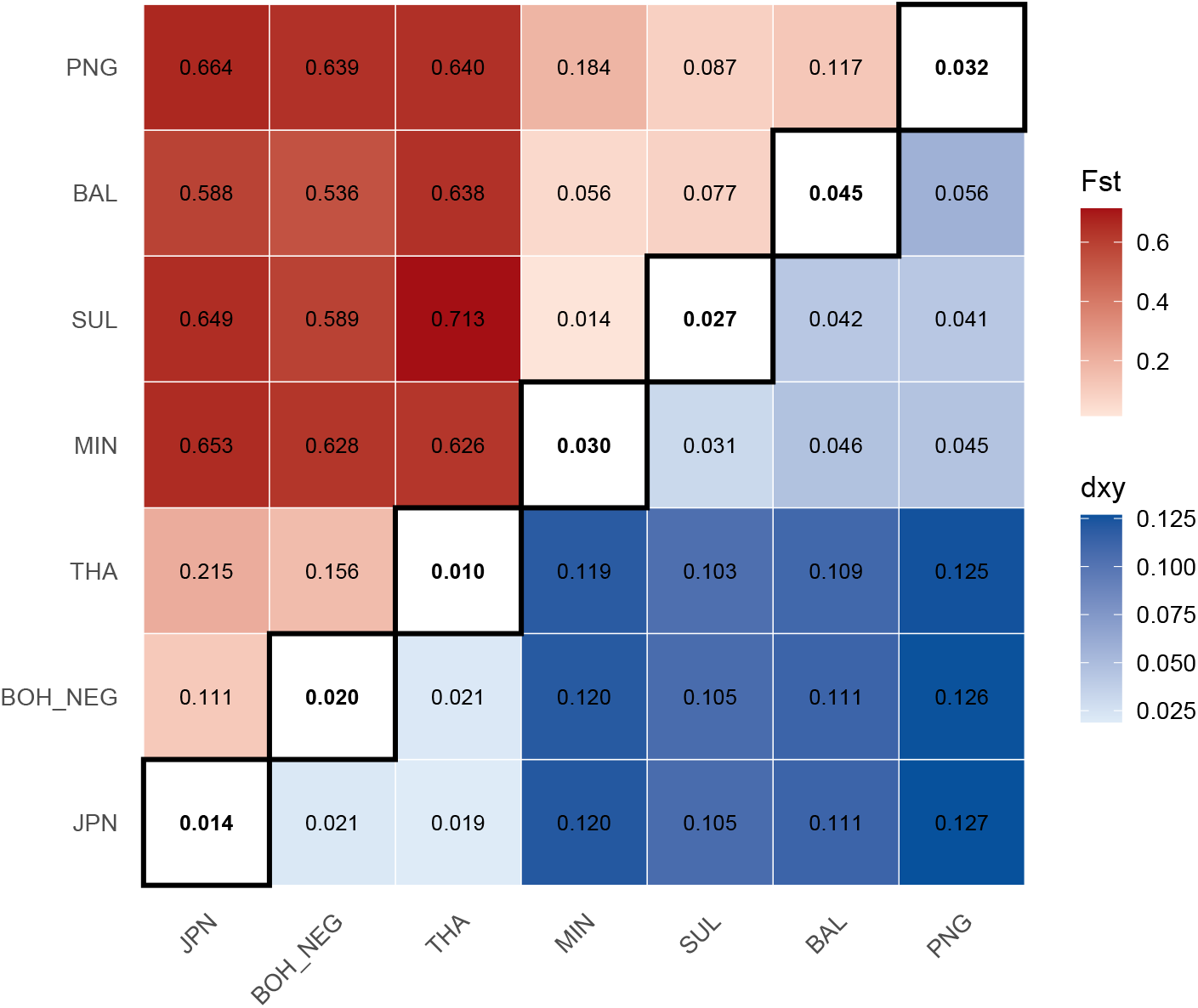
Pairwise *F*_*ST*_ (upper left) and *d*_*xy*_(bottom right) indicate extremely high differentiation between morphs *A. polymnus* (BAL, SUL, MIN) and *A. annamensis* (JPN, BOL/NEG, THA) (*F*_*ST*_ 0.54–0.71), consistent with species-level divergence, whereas *A. laticlavius* (PNG) shows much lower differentiation from *A. polymnus* (*F*_*ST*_ 0.09–0.18) and higher differentiation from *A. annamensis* (*F*_*ST*_ 0.64–0.66). *π* values are shown along the diagonal.

When we examined chromosome 1 (Figure 3), we found an unusual pattern where there is a sharp “peak” in *π* that is 10 Mb wide for all populations except JPN and SUL (Figure 3A). Interestingly, when comparing between populations BOH/NEG vs. all for *F*_*ST*_ values, the average across the chromosome is overall quite high except there is sharply decrease in *F*_*ST*_ between the comparisons of BOH/NEG vs. BAL, MIN, PNG and SUL but not between BOH/NEG vs. JPN or THA (Figure 3B). For *d*_*xy*_, the pattern is opposite of *F*_*ST*_, the overall average *d*_*xy*_ is high across the chromosome for BOH/NEG vs. BAL, MIN, PNG and SUL but there is a small peak 10 mb wide. The comparisons between BOH/NEG vs. JPN or THA are even more striking: overall low average *d*_*xy*_ across the genome with a very high peak in the same location across the genome (Figure 3C).

**Figure 3:**
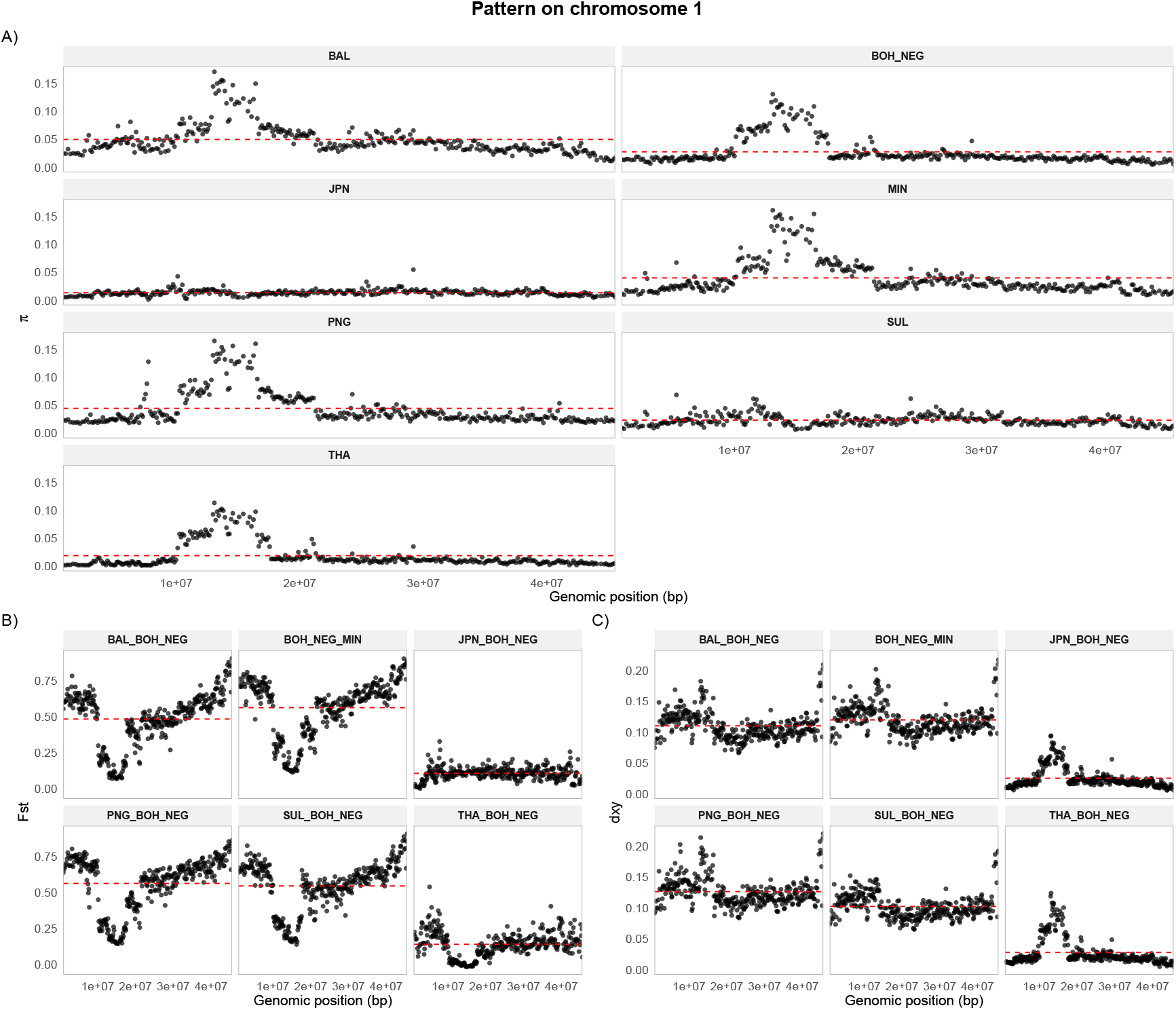
Pattern on chromosome 1 A) *π* values for each population across the chromosome B) *F*_*ST*_ values for BOH/NEG vs. all other populations C) *d*_*xy*_ values for BOH/NEG vs. all other populations. Red line highlights the average values for each comparison across the genome.

### Image Analysis

Using the standardized images dataset, we tested if there were significant differences color pattern based on the morph and found strong differences in PC 1 (variance 28.7%) based on the presence or absence of the full saddle/second vertical bar with the *A. annamensis* morph (small saddle) clustering tightly compared to the *A. polymnus* morph which was more variable in terms of width of the saddle/second vertical bar but still significantly different then the *A. laticlavius* which tend to have differently shaped saddle and orange abdomen (Figure 4). The PERMANOVA test supported the significant difference between all three groups (Figure S1).

**Figure 4:**
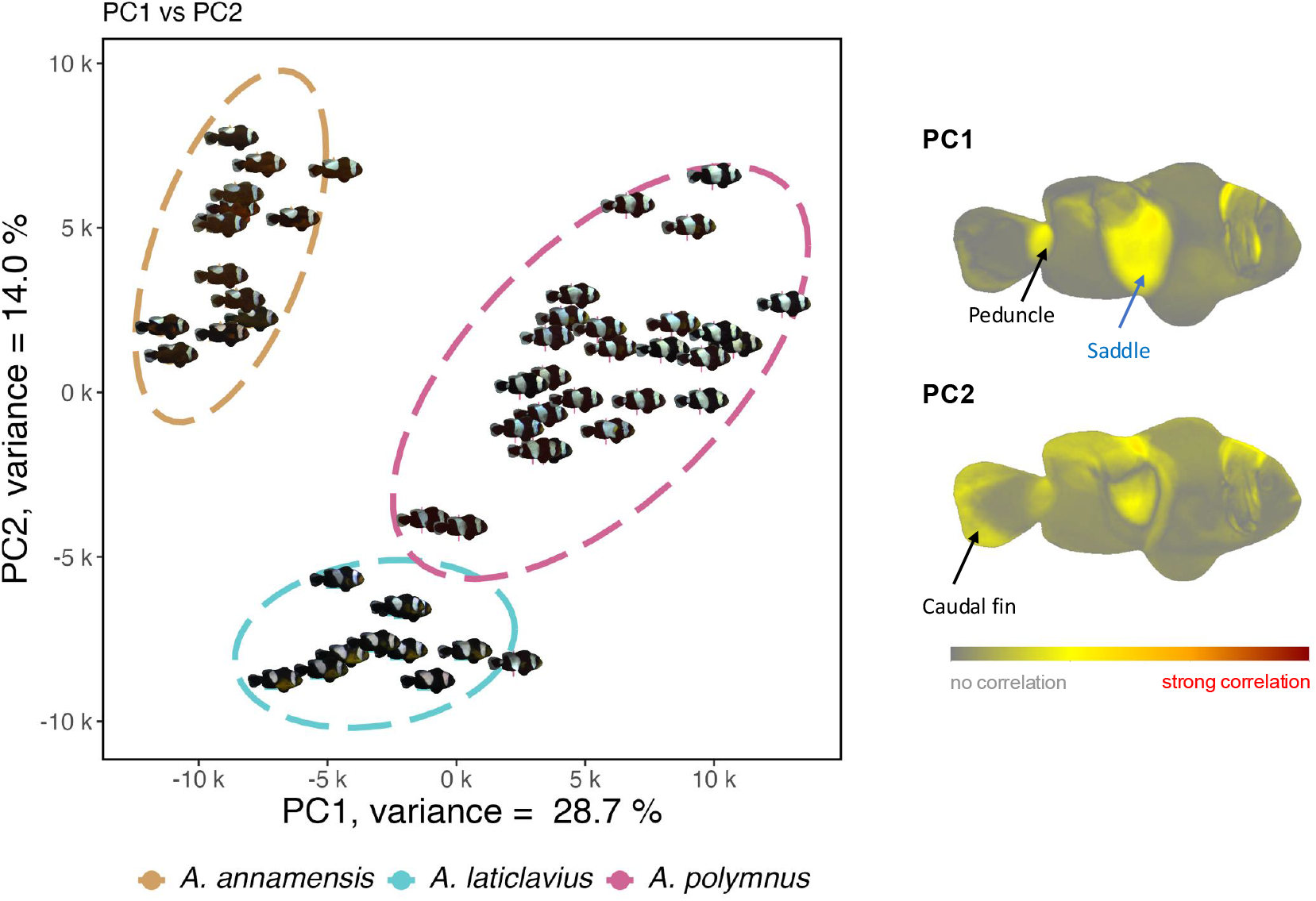
Standardized images (n = 51) PCA showing strong differentiation between the three morphs, circled by color, tan = *A. annamensis*, blue = *A. laticlavius*, and pink = *A. polymnus* with the heatmaps on the right of the figure highlighting which pixels explained the variation on PC1 (upper) and PC2 (lower).

## Discussion

We found strong evidence that color pattern and genetic divergence were associated with the morphs rather than geography. This addresses the central question of whether the observed variation represents intraspecific color polymorphism or divergence among the lineages. Instead of a single polymorphic species, our results suggest at least two genetically distinct groups, one of which may be a species complex.

Genomic analyses revealed clear differentiation between the *A. annamensis* morph from *A. polymnus* and *A. laticlavius* group. ADMIXTURE, supported three ancestries corresponding to the different morphs, while the *F*_*ST*_ and *d*_*xy*_ showed strong divergence between *A. annamensis* and the other two morphs, *A. polymnus* and *A. laticlavius*, but low divergence between *A. polymnus* and *A. laticlavius*. The maximum *F*_*ST*_ value between Sulawesi and Thailand far exceeded the maximum value between *A. clarkii* populations between Maldives and Papua New Guinea (Schmid et al., 2024). This is a striking difference because Sulawesi and Thailand are geographically much closer than Maldives and Papua New Guinea which are separate oceans (Indian vs. Indopacific) indicating that divergence is not driven by geographic distance. Additionally, the pattern on chromosome 1 suggests a structural variant, potentially an inversion, contributing to the observed genetic differentiation. The combination of elevated *d*_*xy*_ and reduced *F*_*ST*_ in this region may indicate conserved or constrained loci shared across morph that could be related to its specialist nature, and live in sandy mucky areas away from coral reefs where other clownfish species live.

Image analysis of the color patterns further support the genetic structure. The first principle components of color variation were strongly associated with features such as saddle width and the presence/absence of the third vertical bar which separates *A. annamensis* from the other 2 morphs. The visual separation was supported by a PERMANOVA confirming that the three morphs were structured rather than continuous. The correspondence between phenotype and genotype contrasts with patterns observed in *A. clarkii* where the coloration did not overlap based on geography and the highest variation was found in the center of the distribution (Schmid et al., 2024). This suggests that in *A. polymnus*, coloration is not plastic or environmentally driven but linked to underlying genetic differentiation.

These findings support the idea that color polymorphism and species complexes can represent different stages along a continuum of divergence. In contrast to the African cichlid example, we did not find clear evidence that the morphs of *A. polymnus* did not occupy different ecological niches as they share similar habitats and host anemone species. The observed genetic and phenotypic differentiation is not driven by ecological divergence.

One explanation could be that the structural variation, potentially inversion, may contribute to maintaining the divergence in the absence of strong ecological partitioning. By reducing recombination, variants can preserve co-adapted regions and facilitate differentiation even when lineages co-exist in similar environments. At the same time, the lack of differentiation between *A. polymnus* and *A. laticlavius* indicates that divergence may still be ongoing. This pattern is consistent with early stages of speciation, where phenotypic differences emerge before complete genomic separation is achieved.

From a taxonomic perspective, our results provide strong support for recognizing *A. annamensis* as a distinct species. This is consistent with the taxonomist who originally synonymized these morphs, and has since supported the recognition of *A. annamensis* as a separate species (G.R. Allen, pers. comm.) although this has not yet been formally published. In contrast, the status of *A. laticlavius* remains less clear. Although there is some differentiation in coloration and ancestry, the overall genomic similarity and phylogenetic relationship suggest its remains closely related to *A. polymnus*. This may reflect an intermediate stage of divergence, where phenotypic differentiation precedes full genetic isolation. Importantly, we did not assess morphological traits beyond coloration and formal species delimitation will require integrate taxonomic approaches. In conclusion, our results highlight that has historically been treated as intraspecific polymorphism may instead represent cryptic lineage diversity with implications for both evolutionary inference and conservation.

## Declarations

LMF and NS conceptualized and designed the study. LMF conducted the research, performed the analyses, interpreted the results, and wrote the first draft of the manuscript. FC performed the image analysis. AM contributed to the genomic analyses. Fieldwork expeditions and sample collection were carried out by LMF, FC, TG, AGJ, PTA, MV, FN, and MM. Standardized photo frames were built by FC and KG. FC, AM, TG, and NS contributed manuscript writing and editing. All authors reviewed, revised, and approved the final version of the manuscript.

## Acknowledgments

We would like to thank G. R. Allen and Putu dian Pertiwi for their feedback on the taxonomic perspective. We would also like to thank Sarah Schmid for her comments on this manuscript. We thank the DCSR infrastructure of the University of Lausanne for the computing resources and the USC Department of Biology for their help in obtaining permits.

## Data Accessibility Statement

Data and scripts will be available on Github. Genomic data in the form of raw reads will be deposited in the NCBI Sequence Read Archive (SRA) under the bioproject ID:XX. Access to these data will also be provided upon acceptance of the manuscript.

## Benefits-Sharing Statement

The sharing of our data and results on public databases ensures transparency and facilitates further research in the field. All fieldwork conducted during this study adhered to local regulations and was carried out in collaboration with local entities. Our research activities were conducted in accordance with the “Access and Benefit Sharing” (ABS) principles of the Nagoya Protocol established by the Convention on Biological Diversity (CBD).

Samples from the Philippines were obtained by Nicolas Salamin’s research group in collaboration with the University of San Carlos (PTA, AD, and PJG contributed to obtaining permits, sampling, and DNA extraction). Samples from Papua New Guinea were obtained by Theresa Rueger’s research group (TR, MV, FN, and KG). Samples from Japan were obtained by Vincent Laudet’s research group (MM and VL).

## Supplementary Figure

**Figure S1:**
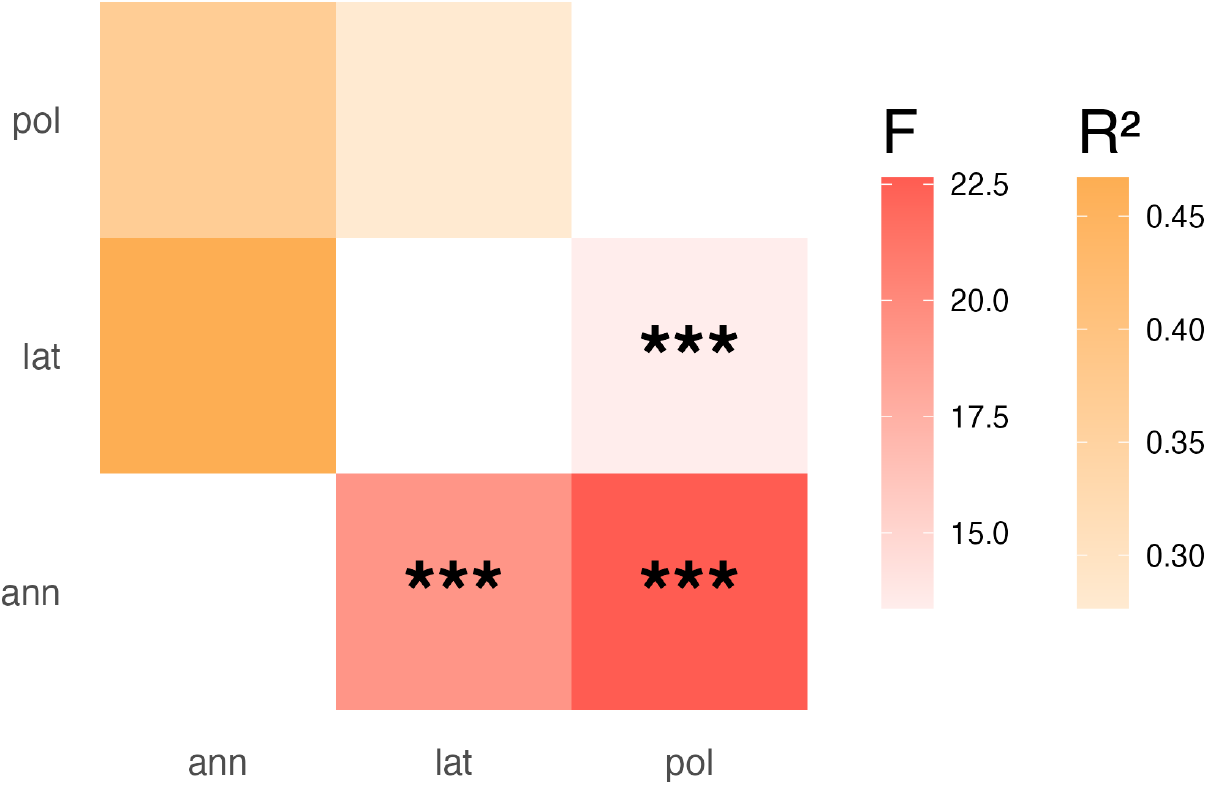
Pairwise PERMANOVA tests quantify the degree of multivariate phenotypic divergence among the three morphs (ann = *A. annamensis*, lat = *A. laticlavius*, and pol = *A. polymnus*. Cells show the corresponding *F* -statistics (red scale) and R2 values (orange scale), with darker colors indicating stronger effect sizes. Asterisks denote significance levels (*** p *<*0.001).

